# Prediction of pH-sensing histidine residues in human cells

**DOI:** 10.64898/2026.09.14.751505

**Authors:** Shalaw Sallah, Jim Warwicker

## Abstract

Changes in pH play an important role in human physiology and cell behaviours, evidenced by studies of tissues and gene expression, and the biophysics of individual systems. Bridging between these scales to establish pH-sensing networks has only been partially explored. Here, amino acid conservation and structural context, and benchmarking with known pH-sensors, were used to predict histidine residues within the human proteome that could modulate pH-dependence. Proteins containing these histidines are enriched in many categories, most notably nucleic acid-binding and transcriptional regulation. DNA phosphate backbone interactions are proposed to play a major role in modulating some pH-sensors. Transcription factors with potential pH-sensors and the largest number of documented target genes are highlighted, including those involved in circadian and developmental processes. Certain CDK-cyclin pairs also feature, along with components of SWI/SNF and cohesin complexes. Focussing on histidines, including those that are solvent accessible in AlphaFold protomer models, this study generates benchmarked predictions for possible pH-sensors in the human proteome.

## 1. Introduction

The control and maintenance of pH is essential in many biological systems. Examples include blood pH, mediated by channels, transporters, and sensors [1], and cytosolic pH [2]. In mitochondria and across the plasma membrane, pH gradients drive key elements of metabolic energy transduction, including proton pathways in ATP synthase [3]. More complex cell types compartmentalise macromolecules and small molecules, including protons, in membrane-bound subcellular organelles [4]. Acidification gradients in eukaryotic import and export pathways support processes that include receptor-ligand cycling, and protease activation [5]. Several aspects of biological pH-dependence are now understood sufficiently to support redesign for altered function. Early studies showed that charge engineering can modify the pH-dependence of enzyme catalysis [6]. More recent work has used directed evolution to engineer enzymes for biotechnology and synthetic biology applications [7], combining sequence and structural features with machine learning (ML) to predict and engineer enzyme optimum pH [8]. Fluorescent proteins with varied pH-sensitivities are central to measuring intracellular pH [9]. A prominent example of exploiting and optimising existing pH-dependence is the extension of protein therapeutic half-life through pH-dependent recycling of antibodies and serum albumin by the neonatal Fc receptor (FcRn) [10]. Studies that tune pH performance often use high throughput mutagenesis to screen successful designs. Examples include engineering antibody–antigen binding to increase affinity at the acidic extracellular pH of tumour environments [11], and designs aimed specifically at creating pH switches [12]. Design and validation have also been combined to adjust buried histidine environments, using the Rosetta suite of tools, to control conformation and oligomerisation through pH switching [13].

An improved understanding of proteome-wide pH-dependence will benefit our ability to target cellular processes, and supplement methodologies for designing pH-dependent systems in biotechnology and synthetic biology. Experimental studies often focus on histidine, with the imidazole sidechain pKa relatively close to physiological pH. A buried and unsolvated histidine in a mildly acidic pH environment is destabilising, an example of electrostatic frustration and a source of pH-dependence [14, 15], an idea also applied to viral protein structure [16]. Histidine can also contribute to pH-dependent effects when stabilised, and with an elevated pKa [17]. Whilst histidine is prominent in pH-dependence, including the active sites of many enzymes, so are other residues, including aspartic and glutamic acids. Networks of proton titratable groups are key features in many enzymes, in acid/base catalysis and proton transfer and in transporters and channels [18, 19]. In recent work, pka calculations and residue burial have been used to predicted charge networks across the human proteome, using AlphaFold protomer models [20]. The amino acids (AAs) identified were named as buried charges of interest (BCOI), recognising that the human interactome (relying on AAs at protomer surfaces) was excluded. Prediction of proteome-wide protein-protein and other interfaces is rapidly advancing, although the co-evolutionary signals need to be enhanced for the complex human proteome and there remains an issue of balancing precision with the number of interacting pairs predicted [21]. The current lack of a definitive human interactome is an issue for attempts to predict pH-dependence from structure-based pKa calculations. A further problem is the accuracy of pKa predictions, which will necessarily be higher throughput than some protocols, for example constant-pH molecular dynamics [22]. Prediction schemes for continuum-based methods typically have a pKa error of at least 1 [23], while ML methods are advancing, with a pKa error of 0.8 in one example [24]. This extent of prediction error matches the range of physiological intracellular pH (pHi) variation (perhaps 6.8 to 7.6 at most [25]). The issues, of interactome interface modelling and pKa prediction accuracy, lead to a requirement for developing alternative protocols to predict proteome-wide pH-dependence.

The AlphaMissense score [26] was found to correlate with predicted BCOI [20]. In the current work two derivatives of the AlphaMissense score have been combined with a separate measure, to which AA conservation also contributes, Mutation Assessor [27]. Thresholds are then developed for best predicting BCOI [20] and a benchmark set of pH-dependent systems, and applied to the human proteome. Histidine is the focus from proton titratable AAs since Asp, Glu, Lys, Arg are much more likely to be involved in salt-bridge interactions that could lead to AA conservation without pH-dependence. The method is a benchmarked and proteome-wide re-working of the process that many laboratories have followed in a single system, identifying conserved His in a search for pH-sensors. Here, application to the human proteome leads to prediction of extensive histidines of interest (termed HOI), that could be pH-sensing. Transcriptional processes are prominent, consistent with experimental work on transcription factors (TFs) [17] and the extent of transcriptional response to change in pHi [28, 29].

## 2. Materials and Methods

Two scores of predicted tolerability to mutation at sites in the human proteome were used to find thresholds that separate histidines identified computationally as members of the BCOI set [20], and identify benchmark cases of known pH-sensing His. AlphaMissense has been developed to predict AA variant pathogenicity using structural context and evolutionary conservation, giving a score from 0 (most benign) to 1 (most pathogenic) [26]. Since AlphaMissense (AM) supplies scores for change to each of the alternative 19 AAs, two derivative scores were made. AM-avg is the average score at a histidine location, over those 19 target AAs. AM-base uses a single score, for change to Tyr, being a close AA in terms of size, hydrogen-bonding and the potential for pi-pi stacking interactions. A third score is added from Mutation Assessor (MA), which gives a functional impact score that reflects amino acid conservation [27]. MA ranked highly in a test of 11 tools for predicting pathogenicity [30], and gives a range of about-4 to 6 (higher being greater predicted functional impact). For context when looking at separation in ROC plots, calculations were also made for AAs normally charged at neutral pH, Asp, Glu, Lys, Arg. In these cases, comparator AAs for the AM-base derived score are Asp/Asn, Glu/Gln, Lys/Met, Arg/Met. The original calculation of BCOI used 10 charge calculation and burial filters, 3 of which were composites from the other 7 [20]. Gene ontology analysis then looked for consistency of BCOI prediction across the calculations [20]. In the current work a clear assessment of BCOI is needed for ROC and distribution analysis, which was made with at least 4 of the 7 non-composite calculations yielding a BCOI prediction.

Enrichment analysis for gene ontology (GO) used the GO knowledgebase [31] and PANTHER and AmiGO tools [32, 33]. Results were interpreted as fold enrichment for the query list relative to the reference list. Code was written to combine GO results for different calculations and to illustrate the GO hierarchy with indentations of the class on each row. The different calculations included separation according to whether proteins include predicted HOI that are accessible or buried or a mixture (solvent accessible surface area, SASA, threshold for accessibility 20 Å^2^). In addition, proteins with HOI were ranked according to the density of HOI (number of HOI divided by length of protein), and a top sliced subset used as the query list in GO analysis. Additional restraints were applied to GO analysis, in most cases to avoid enrichment for classes of protein known to contain functional His, but which are not the focus of this study. The reference list is adjusted accordingly, for example if proteins known to contain zinc finger (ZF) domains are excluded from the HOI query list, then they are also excluded from the human proteome reference set. Annotation of enzyme, transporter, ZF-containing, and ion-binding for human proteins was made from UniProt [34].

Protein and protein-nucleic acid structures were obtained from the PDB [35], with mapping between proteins in structures and UniProt made using the SIFTS data [36] provided by the European Bioinformatics Institute (EBI). A small pipeline was written to process SASA (calculated with local code sacalc) changes in TF-DNA complexes. Swiss PDB Viewer [37] was used for viewing structures and ChimeraX [38] for viewing and figure drawing. AlphaFold2 protomer models [39] were used for SASA calculations on 20503 proteins from the human proteome (UniProt [34]). Predictions of pKa for selected structures from the PDB were made with pkcalc [40], a method used previously in a study of predicted pH-dependence and somatic mutations in cancer [41], and in the BCOI study [20].

Analysis of cyclin-dependent kinase (CDK) – cyclin pairs was assisted by a published alignment [42] and used Clustal Omega [43] at the EBI [44], while CDK-cyclin pairing was derived from the literature [45]. In a study of TF-target pairs, a dataset of TFs [46] was combined with a set of genes whose transcription was monitored when pHi was raised or lowered [29], and with a dataset of measured relationships between TFs and target genes that was constructed using TFLink, TRRUST, and ARCHS4 data [47–49].

## 3. Results and Discussion

### 3.1 Deriving threshold scores for histidines of interest

AlphaMissense and Mutation Assessor, along with benchmarks from a previous study [20] were used to obtain thresholds for histidines of interest (HOI). AlphaMissense and Mutation Assessor give values for mutation to a specified target amino acid (AA), both closely related to AA conservation and in the case of AlphaMissense adding a structural element [26, 27]. Mutation assessor (MA) scores used here are averaged over values for the amino acid changes listed by MA. AlphaMissense (AM) gives scores for all 19 missense changes. The average of these (AM-avg) is one of two AM-derived distributions. The second takes just the single change for each His, to Tyr, more similar in structure to His than other AAs (AM-base). Distributions of MA, AM-avg and AM-base for His AAs are divided into those predicted as BCOI or not [20], and separately into those with more or less than 20 Å^2^ SASA in the AlphaFold protomer models. The aim is to develop thresholds that largely include BCOI AAs, rather than exclude all non-BCOI. Conservation-based metrics should capture AAs of interest that are not buried in AlphaFold model protomers, often due to unseen interactions with protein, nucleic acid, and other ligands. For MA the distribution of BCOI AAs largely follows that of bur20 AAs, and is relatively broad at higher percentages above MA = 2.5 (Fig 1A). For AM-avg (Fig 1B) and AM-base (Fig 1C) distributions, BCOI and bur20 distributions again track each other, but now the distributions are partly bimodal with extensive mid-range AlphaMissense scores that are largely unpopulated. Throughout MA, AM-avg and AM-base distributions there are moderate populations of AAs not recorded as BCOI that score highly. The aim is to set thresholds that maintain a high fraction of BCOI AAs, whilst including also high scoring non-BCOI AAs.

**Fig 1.**
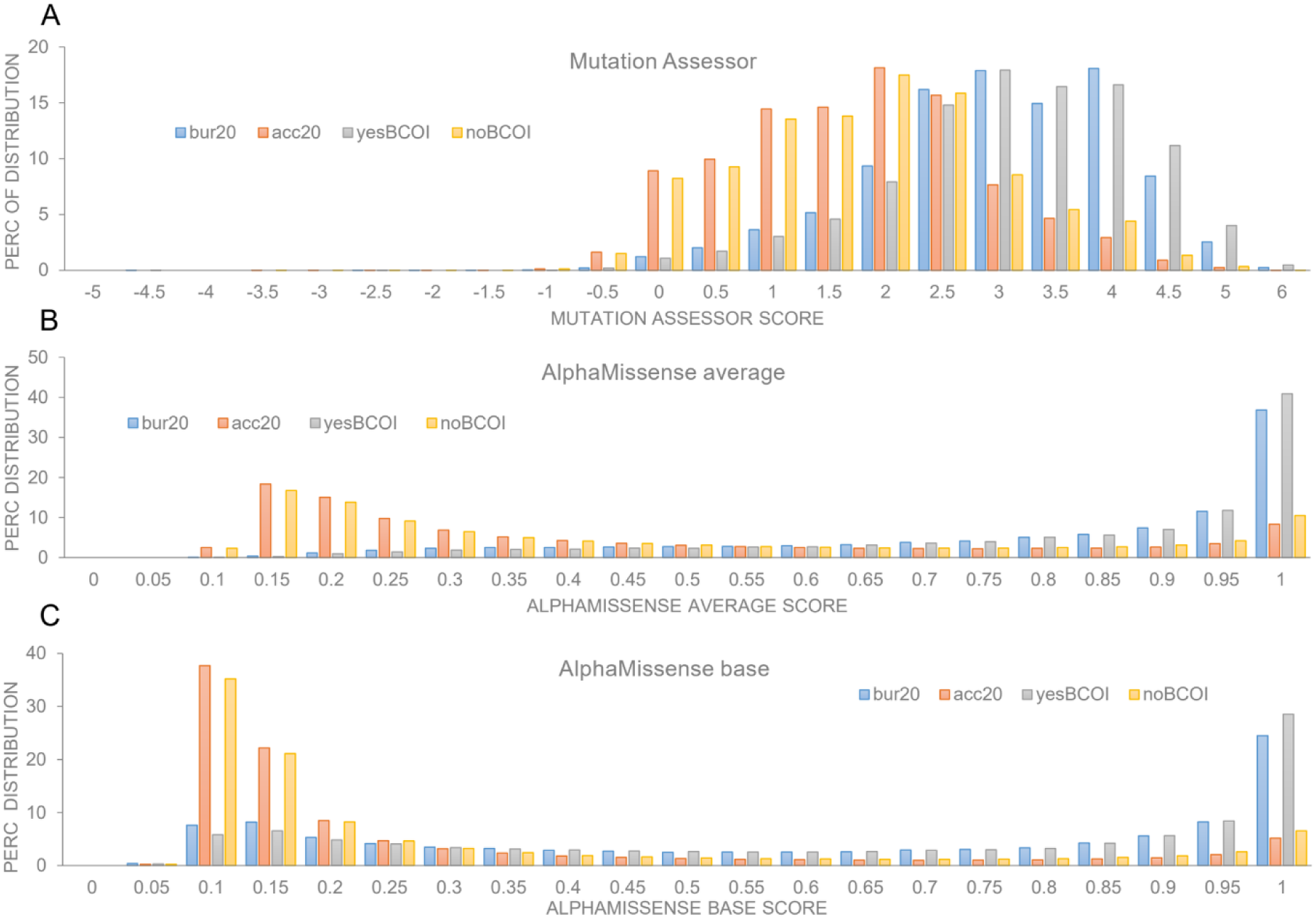
Mutation Assessor and AlphaMissense partition histidines classified as BCOI. (A) Distributions of the Mutation Assessor score are shown for His in the human proteome, separated according to those predicted or not as BCOI, from BCOI assignments for at least 4 of the 7 unique calculation filters used in previous work [20]. Also shown are distributions of His across Mutation Assessor score, on either side of a burial threshold of 20 Å^2^ (bur20 and acc20). (B) Equivalent distributions to those of (A), but with AlphaMissense score averaged over mutation score to other AAs (AM-avg), and (C) with AlphaMissense scores for mutation to a base AA (Tyr) that is closest to His in structure and polarity and not normally charged at or near cytoplasmic pH (AM-base).

Another illustration of separating BCOI from non-BCOI with MA and AM scales is shown in ROC plots (Fig 2). The plots are similar for charge-bearing amino acids Asp, Glu, Lys and Arg, and His. For His AUC values are 0.837 (MA), 0.834 (AM-avg), and 0.828 (AM-base). Although AAs other than His can be involved in pH-dependent function (largely Asp, Glu and Lys, also Tyr and Cys), each of these present extra considerations to His, including conservation for salt-bridge interactions and disulphide formation. For this reason, the current study focusses on His, which is known to play a role in many pH-dependent processes [50–52]. Thresholds to estimate a degree of evolutionary and structural importance that could underpin pH-dependence for His are assessed through a combination of the data shown in Figures 1 and 2, capturing the high BCOI distribution regions of Fig 1, whilst maintaining some non-BCOI (false positive inclusion) but not overwhelming BCOI positive numbers with non-BCOI (Fig 2). The resulting condition for prediction of a histidine of interest (HOI) is MA ≥ 4 or AM-avg ≥ 0.97 or AM-base ≥ 0.97. This returns 32,226 His from 298,836 in the proteome, of which 9,079 are also BCOI, leaving 23,147 non-BCOI His, a similar number to the 23,692 BCOI total His [20]. The chosen thresholds relate to TPR values of 0.301 (MA), 0.457 (AM-avg), and 0.338 (AM-base).

**Fig 2.**
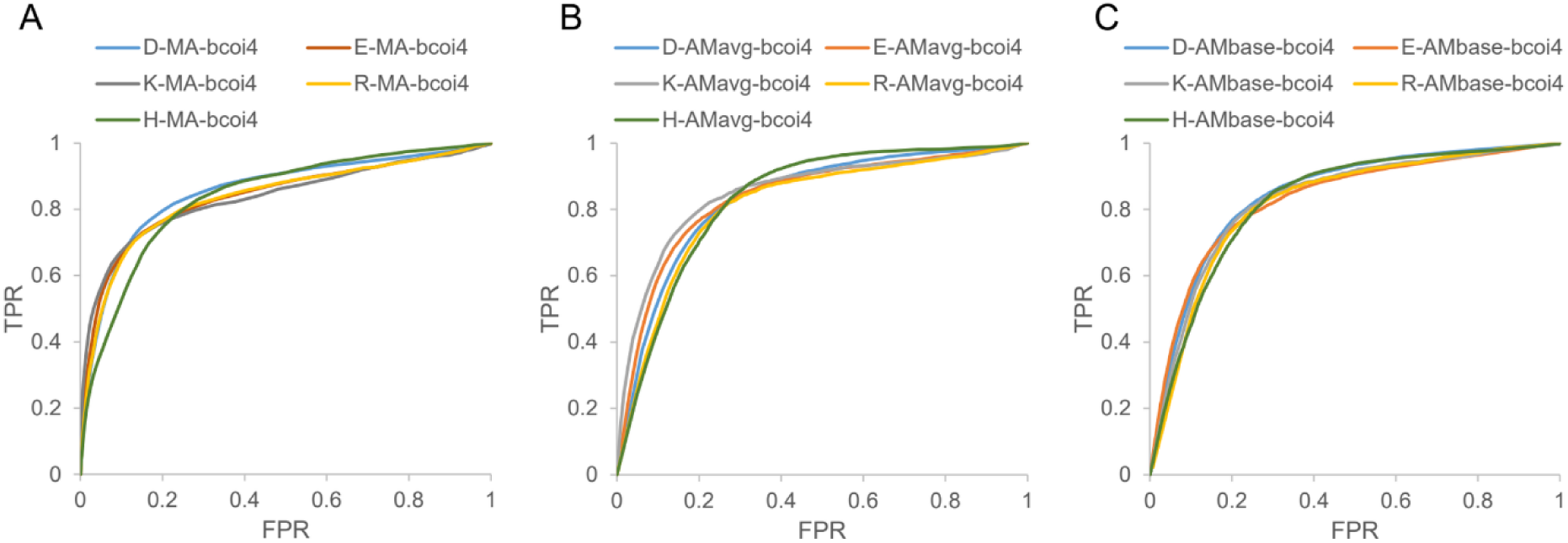
AlphaMissense and Mutation Assessor are comparable in their ability to separate predicted BCOI from non-BCOI. (A) For each of D, E, K, R, and H, ROC plots are shown matching or not predicted BCOI assignments from at least 4 of the 7 unique filters used in previous work (labelled bcoi4) [20], when taking different thresholds from the Mutation Assessor tool (101 points sampled evenly over the score range of-4 to +6). (B) ROC plots as for panel B, but with the AlphaMissense tool, and mutation scores averaged over mutations to all other AAs. (C) Here the ROC plots derive from AlphaMissense scores for the single AA closest in size and polarity, but lacking ionisable group charge that are likely to be ionised close to neutral pH (D/N, E/Q, K/M, R/M, H/Y).

### 3.2 Model performance for known pH-sensor systems

Where a pH-dependent system has been discovered, researchers will typically look at sequence conservation and focus on histidine, to narrow down mutagenesis studies that could establish a pH-sensor. The current prediction scheme is a focussed and benchmarked version of this methodology. In previous benchmark testing for prediction of pH-dependence via BCOI [20], BCOI groups were recorded for 11 of 15 systems, with 26 groups reported to be pH-sensing AAs in these systems and 16 were predicted as BCOI. In the current work, using MA and AM thresholds, 10 systems have predicted HOI. Several within that set of pH-sensing proteins, particularly channels and transporters, have reported Asp and Glu pH-sensors, although these may also be in charge networks that include His. To focus a benchmark test on His pH-sensors, the recent peer-reviewed literature was searched for examples where pH-dependence had been localised to a specific histidine by mutagenesis. Ten systems are listed in Table 1, all of which contain HOI (in comparison with the 9,006 of 20,503, 43.9%, proteins in total with HOI).

**Table 1.**
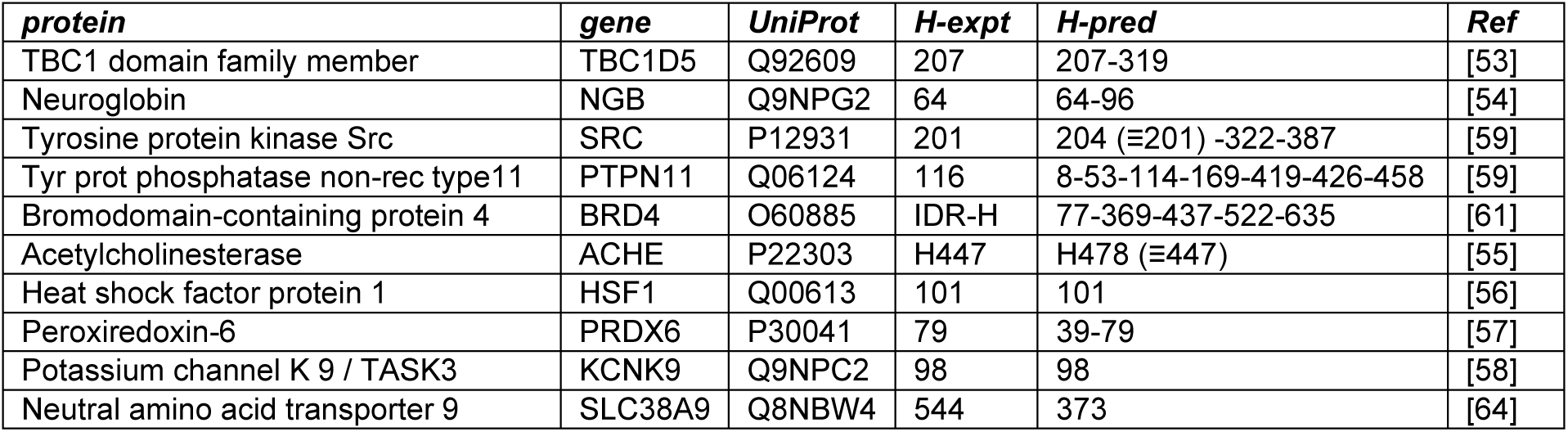
Reported pH-sensing histidines compared with predicted HOI in 10 systems.

For 7 of the 10 systems the identified His sensor is reproduced. Those 7 systems cover a range of activities. TBC1D5 is a GTPase-activating protein that imparts a pH-sensing activity to endosome maturation [53]. Neuroglobin is a brain-localised oxygen storage and transport protein, with H64 regulating ligand binding at the distal heme location [54]. Acetylcholinesterase breaks down the neurotransmitter acetylcholine, with catalytic His H447 also sensitive to pH-dependent methylation [55]. Trimerisation of HSF1 is required for DNA binding by this master transcriptional regulator of heat shock factor expression, and is sensitive to pH via H101 [56]. A pH-dependent oligomerisation of peroxiredoxin-6 modulates activity, with D42 and H79 the key amino acids [57]. Potassium ion conductance of TASK-3 contributes to modulation of membrane potential and cell excitability. H98 and a surrounding hydrogen-bonding network control the pH-dependence of TASK-3 activity [58].

One of the 3 systems where specific HOI is not predicted (SHP2 phosphatase) was reported alongside a pH-sensitivity of the Src tyr kinase, for which the correct HOI (H201, [59]) is identified. For SHP2, a pH-dependent interaction between H116 and E252, that modulates activity, was inferred from mutagenesis. Predicted HOI include H114 but not H116, which is clearly below the thresholds for HOI prediction (e.g. AM-avg of 0.804 compared with the 0.97 threshold). Although the current work does not pursue a model for AAs other than His, E252 is substantially above the threshold used for HOI (AM-avg of 0.997). Interestingly, the SH2 domain H201 of Src maps to SH2 domain H169 of SHP2, also an HOI. A recent deep mutational scanning analysis of SHP2 identifies H114, a predicted HOI, as involved with regulation of phosphatase activity [60]. Nevertheless, SHP2 H116 has been reported as a functional pH-sensing AA but was not identified in the current work.

None of the predicted HOI for BRD4 locate to the intrinsically disordered regions (IDRs) that are shown to add pH-dependent control to the formation of transcriptional condensates [61]. This highlights an interesting issue since IDRs are generally less conserved than structured regions that form a specific fold [62]. A further complication is that enrichment for AA type, rather than conservation at a single site, will mediate IDR properties such as pH-dependence. None of the BRD4 IDR His are close to the HOI thresholds of the current study, showing that there is scope for improvement when considering pH-dependence mediated by IDRs, perhaps lowering the thresholds via benchmark testing of IDRs and considering AA enrichment rather than single sites. It is possible that pH-dependence for BRD4 is not restricted to IDRs. Predicted HOI H77 is adjacent to one of the basic patches reported to interact with nucleosome DNA [63].

Mutating H544 of SLC38A9 abolishes the pH-dependence of arginine uptake [64], whilst the sole predicted HOI is H373. A study of buried charges of interest (BCOI) for the human proteome placed H544 as part of a network (R167-E354-K422-H544) for SLC38A9 [20]. Although prediction schemes overall implicate H544, the specific HOI prediction fails, and H544 is not borderline threshold (e.g. AM-avg = 0.85). In sequence logos examining conserved His [64], that for H544 was the most extreme in terms of higher conservation than neighbouring AAs, suggesting a direction for further investigation. Overall, Table 1 shows that model performance is encouraging.

### 3.3 Cellular components relating to nucleic acid binding are amongst those enriched for proteins with predicted HOI

Certain categories of proteins are known to be enriched for conserved histidines, including enzymes, transporters, and proteins containing zinc-finger (ZF) domains and other ion-binding motifs [20, 65]. Any such proteins could, in principle, exhibit pH-dependent activity through their HOI. Enzymes often have pH optima associated with acid-base catalysis [65], many transporters have been recorded as pH-dependent (OATP1B3 for example [66]), and ion binding sites can balance proton affinity with metal binding affinity [67]. These are known classes of proteins rich in HOI. Since the goal of this work is to highlight lesser-known classes that appear in HOI analysis, unless stated otherwise those known HOI classes are omitted. Removing proteins annotated in UniProt [34] as enzyme, transporter, containing a ZF domain, or with ion-binding properties, reduces the 20,503 background set of human proteins to 13,469. The 9,006 proteins returned as containing HOI are reduced to 5,800 (without enzymes), then 5,685 (without transporters), 4,521 (without zinc finger domains), and finally 4,093 (also without annotated ion binding).

From analysis of fold change and false discovery rate (FDR) with 4,093 HOI proteins against 13,469 background, a-log10(FDR) threshold of 5 was chosen (Fig 3A) for inclusion in a heat map of GO Slim cell component terms (Fig 3B). Subsets of the 4,093 set were added to examine the influence of buried and solvent accessible HOI, and 1,300 proteins with particularly high density of HOI (number of HOI normalised by protein length). Much of the signal arises from proteins that contain solvent accessible HOI, i.e. that would have been missed by analysis of charge burial in AlphaFold protomers [20]. This observation supports use of scores related to AA conservation to identify HOI outside of Alphafold protomer folding cores, many of which are likely to be at interfaces. The density of HOI sites column does not substantially alter the analysis, although it does reinforce results for some GO categories in the depletion map (Fig 3C).

**Fig 3.**
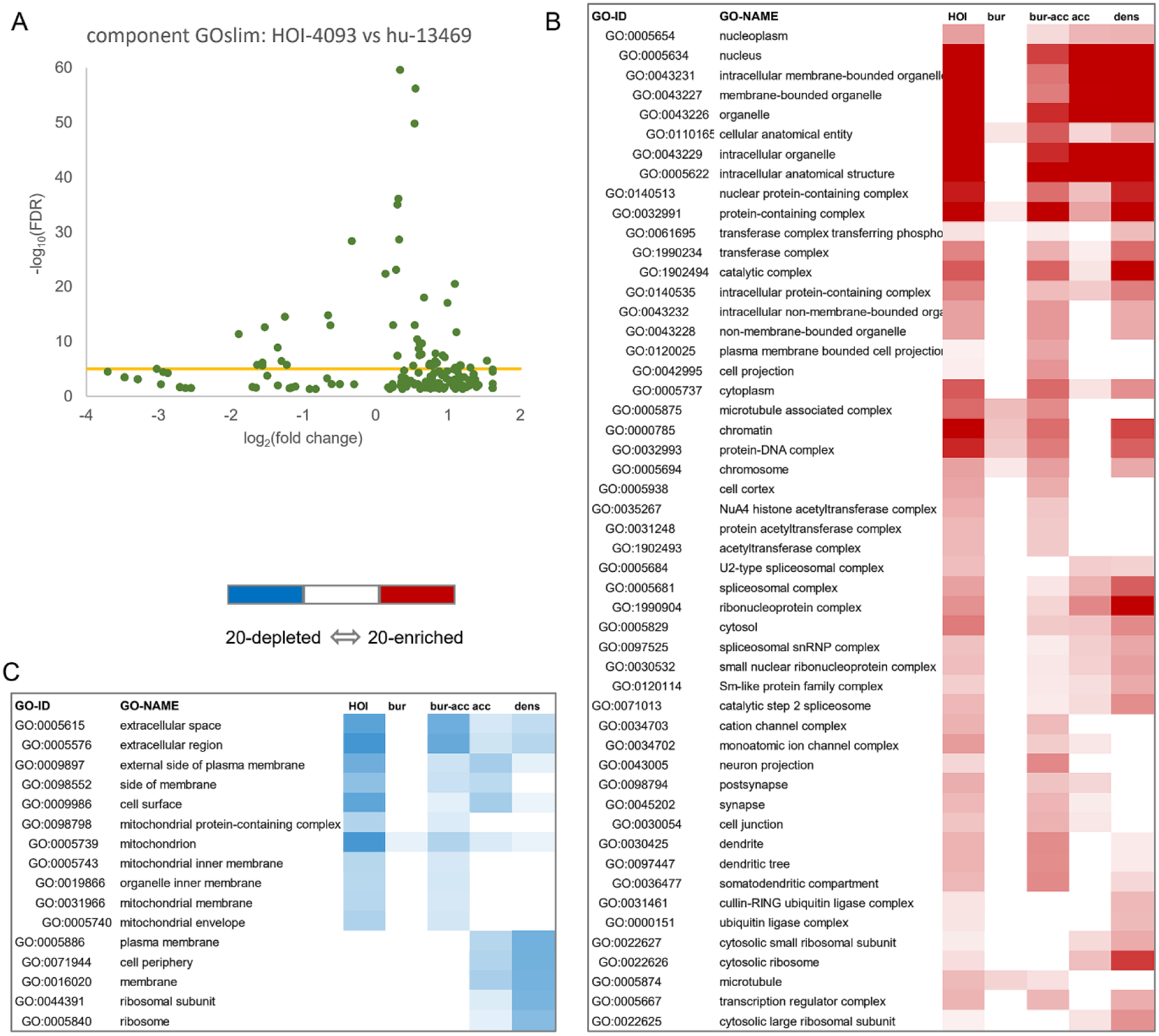
Gene ontology analysis identifies cellular components enriched and depleted in predicted HOI. (A) Volcano plot of GO Slim component terms returned when 4,093 human genes with HOI are examined against a background set of 13,469 genes. A line drawn at-log_10_(FDR) = 5, emphasises that a cut-off based on false discovery rate is more effective than one based on fold change in identifying the most significant enrichments. (B) GO Slim cell component terms enriched in HOI genes are shown where any of 5 calculations give - log_10_(FDR) ≥ 5. GO categories are given by code and name, and nested where child (above) and parent (below) relationships are known. The 5 calculations each give a different emphasis on the data: the 4,093 genes with any HOI, the 1,249 genes with only buried HOI (SASA < 20 Å^2^), 1,189 genes with both buried and accessible HOI, 1,655 genes with only accessible HOI (SASA ≥ 20 Å^2^), and 1,300 genes that rank highest when sorted by density of HOI occurrence. (C) Equivalent data to panel (B), for GO Slim cell component terms that are depleted in predicted HOI genes. Heat map colour colours have thresholds at-log_10_(FDR) = 20 for both panel B (red, enrichment) and panel C (blue, depletion).

GO categories referring to nucleic acid binding are the most consistently enriched for proteins with HOI, for example nucleus, nuclear-containing protein complex, chromatin, ribonucleoprotein complex (Fig 3B). In the smaller list of GO categories that are strongly depleted for proteins with HOI, the mitochondrion and cell surface / extracellular space feature across most of the columns. An equivalent analysis to that for HOI has been made for glutamine, an AA with relatively close sidechain chemical similarity to histidine, but without the sidechain ionisation close to neutral pH. The glutamine heat map is not as extensive as that for HOI (S1 Fig), but similar in enrichment of proteins that bind nucleic acid. His and Gln have similar reported propensities for locating to interfaces with DNA [68]. It is likely that both HOI and conserved Gln are identifying interfaces, although histidine more extensively and with the aspect that makes it of interest, potential pH-dependence.

### 3.4 Transcription factors are enriched in HOI

Noting the prevalence of nucleic acid complex GO categories (Fig 3), TFs were studied more closely. A pH-dependence of TF-DNA binding has been measured for a small number of TFs in a wider set noted to contain conserved histidines [17]. Of 104 His identified by experiment and AA conservation in the FOX, SOX, and KLF families, and a selection of basic helix-loop-helix leucine-zipper (bHLH-LZ) TFs [17], 99 are predicted HOI, with the distribution of AM-avg scores clearly differentiated from those for background His and DEKR (charged AAs) in the same TFs (Fig 4A). Four of the missing HOI are in KLF family members. Alone amongst this set of TFs the KLF family contains ZF domains, specifically included here for comparison. The two alignment positions noted previously [17] are one ZF and one non-ZF HOI. Typically, HOI predictions return His both within and outside the ZF metal ion site.

**Fig 4.**
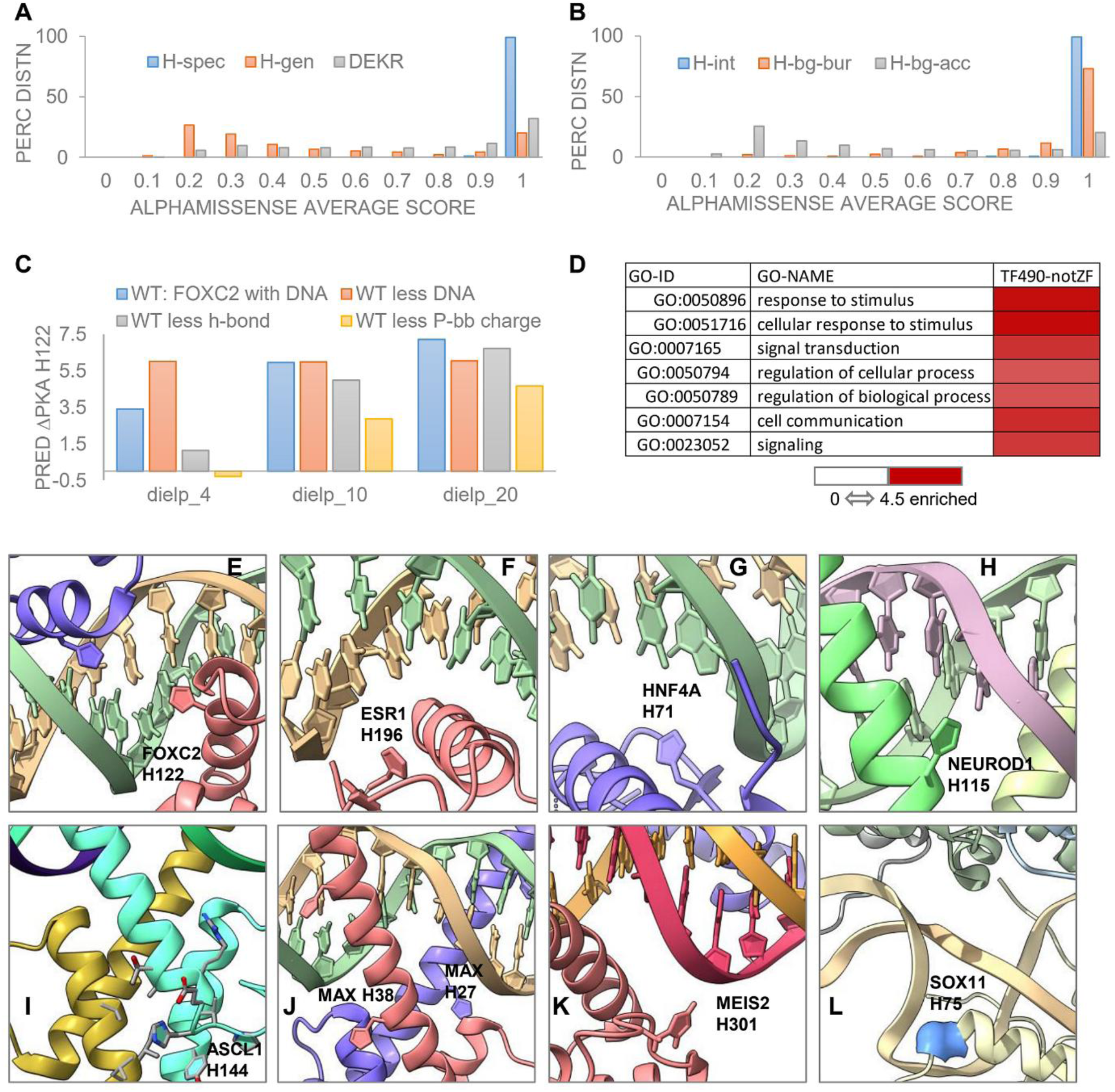
HOI sites identify histidines at protein – DNA interfaces. (A) For the 99 FOX, SOX and KLF TF family members previously identified with conserved His [17], distributions of AM-avg score for the HOI (H-spec) are shown, along with histidines in the same proteins that are not HOI (H-gen), and charged groups (DEKR). (B) For structures of TFs bound to DNA from the PDB [35], with HOI that alter SASA when DNA is removed, AM-avg distributions are shown for those HOI (H-int), and for the remaining His in the same TFs, split according to burial at a threshold of 15 Å^2^ (H-bg-bur) or accessibility (H-bg-acc). These data contain only one structural instance of each HOI. (C) Various pKa predictions (pkcalc [40]) were made for HOI and known pH-dependent AA H122 in FOXC2 [17] bound to DNA (6ako [77]). Three relative dielectric values were used for solvent excluded volume, 4, 10, 20, as indicated. Systems studied were the native FOXC2–DNA complex (WT), WT less the hydrogen-bond charge for an interaction between H122 and a DNA base, FOXC2 alone, and the WT complex but without charge on the phosphate backbone. (D) From a dataset of 1,639 human TFs [46], 1,296 have predicted HOI. Excluding those known to contain ZF domains gives a query set for GO Slim process analysis of 490 TFs with HOI against a background set of 764 TFs without ZF domains. Some prominent enrichments with-log_10_(FDR) ≥ 3 are shown. Panels (E) to (L) show examples of HOI adjacent to either the interface with DNA or a PPI (TF dimer interface). (E) FOXC2-H122 dimer copies, Forkhead family, 6o3t [78]. (F) ESR1-H196, nuclear receptor, 1hcq [79]. (G) HNF4A-H71, nuclear receptor, 3cbb [80]. (H) NEUROD1-H115, bHLH, 2ql2, human numbering equivalent from mouse structure [81]. (I) ASCL1-H144 (at TF dimer interface), bHLH pioneer TF bound to nucleosome, 8ubk [82]. (J) MAX-H27 (DNA interface) H38 (dimer interface), bHLH-LZ, 1hlo [83]. (K) MEIS2-H301, TALE/homeobox, 4xrm [84]. (L) SOX11-H75 (mainchain only shown, adjacent to nucleosome DNA), HMG box pioneer TF, 6t7a [85].

To examine HOI at known TF-DNA interfaces, the RCSB of August 2025 was queried for coordinate files containing human TFs (20,221). A set of HOI (13,803) that excludes enzymes and transporters but includes ZF-containing proteins was derived with the added condition of an AlphaFold protomer model SASA ≥15 Å^2^ (to focus on interface AAs). The UniProt IDs of the HOI-containing proteins was matched to pdb codes with the EBI SIFTS tool [36], with 4,221 redundant mappings for the HOI. Considering only HOI where DNA is present in a coordinate file, and with a change in HOI SASA when DNA is removed, leaves 882 sites and 244 when made non-redundant. These HOI are then termed interacting (with DNA), and other His in the final set of TF-DNA complexes are treated as a background, which is further divided into those accessible (1,872 His) or buried (230 His) in AlphaFold protomers. DNA-interacting HOI exceed the AM-avg score of buried His, some of which are likely to be zinc binding, and far exceed that of background accessible His (Fig 4B). This analysis of TF-DNA complexes supports the bioinformatics identification of HOI.

An important question is to what degree HOI are viable predictions of sites that could modulate pH-dependence around physiological pH. Previous work has demonstrated this is the case for a small subset of TF–DNA interactions, including H122 (a predicted HOI) of FOXC2 [17]. FOXC2 binding to a FkhP DNA motif is modulated between pH 7 (tighter binding) and pH 7.5 (lower binding), with H122K removing the pH-dependence and giving tighter binding, and H122N removing the pH-dependence, with lower binding [17]. Predictions of H122 pKa were made with the pkcalc combined FDPB-DH method [40]. The protein/DNA relative dielectric value was varied between 4 and 20, reflecting commonly used values. Histidine 122 pKa calculated for wild-type (WT) complex only matches the experimental range when the relative dielectric is increased to 20 (predicted pKa = 7.2, Fig 4C). This is consistent with the higher dielectric values used in some FDPB protocols, often where conformational relaxation is excluded in a static calculation [69–71], and with measured values for duplex DNA [72, 73]. Having established that pKa calculations can be used to interpret HOI, the next question is which components in the TF-DNA complex are most important in terms of charge interactions for FOXC2-H122. These are shown in Fig 4C, with the caveat that calculations are made for removal of components from the WT complex in the absence of conformational relaxation. Loss of the hydrogen-bond interaction between H122 and a DNA base gives a small reduction in predicted pKa, with a larger effect for removal of DNA entirely. In the latter case (free protein), the predicted H122 pKa of 6.1 (with a relative protein/DNA dielectric of 20) is close to the model compound value used in these calculations (6.3). Strikingly the pKa shift is much larger (down to 4.7) when the TF-DNA complex is intact, but charges of the phosphate backbone are removed. Phosphate charges form a cradle around TF charge in the major groove, in this case with a predicted ΔpKa contribution of 2.5. This is larger than that for removing DNA entirely, due to the stabilising effect of increasing H122 hydration when DNA is removed. The proposed size of phosphate cradle interaction with protein charges in the major groove (and presumably also in the minor groove) is not unique to His since H122K rescues the higher affinity binding for FOXC2 [17]. Analysis of detailed charge positioning, solvation, net charge balance, and DNA conformation could improve understanding of how Lys and Arg interactions with the phosphate backbone stabilise protein-DNA complexes. What is important for His, but not Lys or Arg, is the pH-dependence around physiological pH. It may be possible for groove-located and destabilised Asp and Glu to have pKa altered to above 6 if the magnitude of the phosphate cradle charge effect is close to that predicted for FOXC2-H122. There is some evidence for acidic sidechain contributions to pH-dependence in protein-DNA complexes [74] although the extent of pH variation was postulated to be lower than for protein-protein interactions due to reduced variation in the environment [75].

Having established that TF-DNA interactions are of particular interest with respect to HOI, the entire set of annotated human TFs was studied. From a dataset of 1,639 human TFs [46], 1,630 matched the 20,503 human proteome of the current work, of which 1,296 contain at least one HOI, and 490 of those are not annotated as containing a ZF domain. Overall, 764 of the 1,630 TFs are not annotated with a ZF domain. GO enrichment was calculated for the set of 490 TFs with an HOI against the 764 set (with and without HOI), all annotated as lacking ZF domains. Whilst this limits the analysis to non-ZF TFs, it avoids the complication of separating HOI inside and outside of ZFs, and the added complication of possible coupled proton and zinc ion sensing. For GO Slim process categories, the most significant enrichments include signalling processes from the cell exterior and the nuclear receptor family (Fig 4D).

Panels E to L of Fig 4 show examples of HOI locations within some of the 490 TFs (Fig 4D). FOXC2-H122 (Fig 4E) shows the major groove location analysed in Fig 4C. Nuclear receptor HOI, ESR1-H196 (estrogen receptor, Fig 4F) and HNF4A-H71 (fatty acid binding, Fig 4G), are adjacent to DNA. A report of pH-dependence for ESR1 identified H196 and E203 as being responsible, with conservation throughout the 50-member nuclear receptor family [76]. It is interesting that pathways relaying signals from outside the cell, with a greater pH variation than inside the cell, also exhibit pH-dependent transcriptional activity in the nucleus. Three TFs containing bHLH domains are shown (Fig 4H-J). NeuroD1-H115 interacts with the phosphate backbone of its target DNA. H144 of the pioneer transcription factor ASCL1 is at the TF dimer interface in a complex bound to nucleosome DNA. For MAX, one HOI (H27) is at the DNA interface and one (H38) at the TF dimer interface. MEIS2 is a Tale/homeobox TF with H301 at the DNA interface (Fig 4K). H75 of another pioneer TF (SOX11 of the HMG-box family) sits at the interface with nucleosome DNA (Fig 4L).

Excluding enzymes, transporters, known ion-binding proteins and those that contain zinc finger domains, the overall fraction returned with HOI (4,093 of 13,469) is 30.4%, but for TFs that doubles to 64.1% (490 of 764). This degree of predicted pH-dependence in TF–DNA interactions is precedented from studies of nuclear receptor and other TF families [17, 76], and from the global changes in transcription have been observed upon modulation of pHi [29]. An experimental pH shift of 0.5 gives about 4 kJ/mole binding change for the FOXC2–DNA interaction [17]. If physiological ΔpHi is around half of this [25], binding energy changes will be in the low kJ/mole, but could contribute a binding constant change of around 2. Current results suggest that small binding changes with ΔpHi could be present in a significant part of the human TF repertoire.

### 3.5 HOI are enriched in some cyclin dependent kinase – cyclin pairs

Given the high proportion of TFs returned as containing HOI, and reported coupling between expression levels, pH and the cell cycle [28, 86–88], HOI analysis was applied to the cell cycle. First, to include CDKs, GO term enrichment was calculated for the 1,300 proteins most dense in HOI in the enzyme and transporter set (4,810 in total). Several GO Slim process categories are enriched for those most dense HOI proteins (Fig 5A). Next, non-enzymes were studied with the same GO analysis used for Fig 3, but extracting categories of greater relevance to the cell cycle and showing enrichment (or depletion) mostly at less significant FDR than terms highlighted in Fig 3 (Fig 5B).

**Fig 5.**
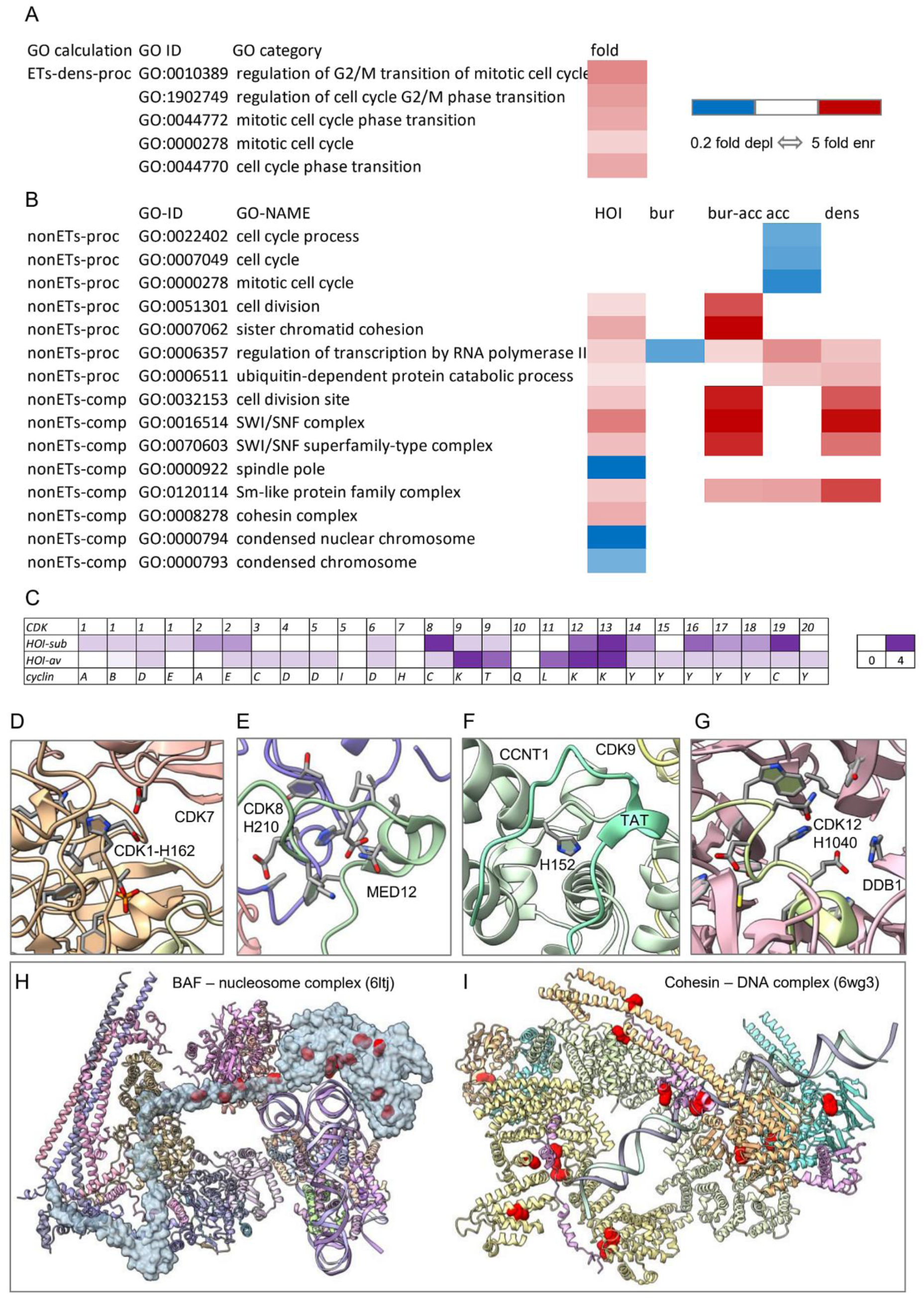
Enrichment of GO categories for proteins with HOI coalesce on particular CDK-cyclin pairs. (A) For enzymes and transporters (ETs, including CDKs), enrichment of GO Slim process terms are shown for the 1,300 proteins most dense in HOI (ETs-dens-proc), against a background set (4,810) of ETs (4,451 enzyme and 383 transporter annotations, with 24 in both). (B) GO Slim process and cell component terms enriched in HOI genes are shown using the same non-ET and non-ZF/ion binding sets as Fig 3 (nonETs-proc and nonETs-comp, 4,093 proteins against 13,649 background, with subsets according to HOI SASA). In both panels A and B fold change is shown (depleted to enriched as indicated in the colour key). (C) CDK – cyclin pairs are shown with the number of HOI associated with each protein. HOI-sub for CDKs indicates subtraction of two HOI that are close to the enzyme active site and not interfacial. For CDK1 these HOI are H120 and H126, although histidines are not normally incorporated into catalytic models for CDKs [95]. For 3 CDKs (10, 11, 20) with zero HOI-sub, both active site adjacent His are present but only 1, 0, 1 are returned as HOI, respectively. HOI-av for cyclins indicates that HOI number is averaged over any multiple sub-types for a listed cyclin. Panels (D) – (G) show examples of HOI in complexes with CDKs and cyclins. (D) CDK1-H162 at an interface with CDK7 (9skq [96]). (E) CDK8-H210 at an interface with MED12 (8tqc [97]). (F) Cyclin T1 H152 is close to an interface with the viral Tat protein (4ogr [98]). (G) CDK12-1040 lies at an interface with DDB1 (6td3 [90]). (H) HOI locations are shown in red on the blue surface of SMARCA4 within a BAF–nucleosome complex resolved at 3.7 Å (6ltj [99]). (I) Locations of HOI (red) in a 5.3 Å resolution structure of a cohesin–DNA complex (6wg3 [100]).

Predicted HOI numbers for CDKs (Fig 5C) have been modified to remove two His that are buried and adjacent to the active site. HOI are associated with most CDK-cyclin pairs but concentrate in some (Fig 5C). Interpretation is limited by the available experimental structures, with 4 examples shown (Fig 5D-G). In CDK1, there is just one HOI distinct from the active site, H162, present also in CDKs 2, 3 and 7. CDK1-H162 is one of 41% HOI in CDKs that are classed as solvent accessible (SASA ≥ 20 Å^2^), where conservation could arise from interfacial properties not recorded by AlphaFold protomer calculations. CDK1-H162 locates to the kinase activation loop close to interfaces in structures, for example as part of a charge network at an interface with CDK7 (Fig 5D). This network (including an aspartate from CDK7) could mediate elevation of the pKa to physiological pH and introduce a pH-dependence of complex formation. Of human CDKs, CDK8 has the most HOI returned (9, Fig 5C). In a structure of the CDK8 kinase module that also includes cyclin C (Fig 5E), CDK8-H210 lies at the interface with MED12 (mediator of RNA polymerase II transcription subunit 12), again with neighbouring acidic charge that could mediate an HOI pKa shift to the physiological range. Additionally, CDK8-H75 lies at an interface with MED13 in the same complex, with neighbouring acidic groups (not shown). Reference to RNA polymerase II, and a potential link to pH-dependent modulation of transcription, is continued with HOI in the P-TEFb (positive transcription elongation factor b) complex that includes CDK9 and cyclin T1 (Fig 5F). T-type cyclins are amongst the highest in terms of HOI, but with variation. Cyclin T2 has 5 HOI and cyclin T1 just one HOI, yet all the 5 HOI predicted for T2 are present in T1. The HOI returned for cyclin T1 (H152) lies close to the interface with viral Tat protein (Fig 5F), as does a cyclin T2 HOI that is just missed in cyclin T1 (H154, AM-avg = 0.951). Separately, a histidine-rich domain of CCNT1 has been reported to modulate binding of CDK9-cyclin T1 to RNA Pol II [89]. Histidines within that domain are relatively low in AM-avg score, suggesting (as for BRD4) that intrinsically disordered regions present additional difficulty in recognising HOI, presumably in part due to lack of intrinsic structure, but also through more distributed activity in groups of AAs. CDK12 and cyclin K are heavily populated with HOI (Fig 5C). In a structure that includes cyclin K, one of the CDK12 HOI (H1040) lies at an interface with DNA damage-binding protein 1 (DDB1) in a charge network that is modulated by burial (Fig 5G). This interesting case is likely to be outside of normal physiological function, since the CDK–DDB1 interaction is promoted by a CDK-inhibitor, part of a screen for small molecules that could promote protein degradation (6td3 [90]). Conservation presumably derives from the role that CDK12-H1040 plays in a C-terminal extension that modulates activity in this kinase [91].

Some proteins related to the analysis of HOI and cell cycle (Fig 5) are highly enriched in HOI. One example is the SMARCA4 enzyme component of the SWI/SNF (BRG1/BRM-associated factor, BAF) remodelling complex, which is also enriched in HOI-containing proteins for non-enzyme components (Fig 5B). For 17 HOI returned within the 1,647 AAs of SMARCA4, about half lie outside of the catalytic domains [92]. In a structure of nucleosome-bound BAF, HOI sites are shown throughout SMARCA4, many proximal to the nucleosome (Fig 5H), suggesting that pH-dependence could play a role in gene expression, including within the cell cycle. Histidines in a low-complexity domain of yeast SWI/SNF component SNF5 mediate pH-sensing that couples to carbon starvation [93]. That region is missing from the human orthologue SMARCB1, and whether pH plays any role in SWI/SNF function, outside of catalytic activity, is unknown. A structure for the cohesin complex contains enzymes and non-enzymes (Fig 5I), the latter contributing to the enrichment for HOI-containing proteins in the cohesin complex (Fig 5B). Non-enzyme cohesin subunit SA-1 is the largest contributor of HOI (7). The large number of cohesin complex HOI suggests that pH modulation in nuclear complexes may extend to DNA replication. A change in cohesion strength when pHi is altered has been reported [94].

### 3.6 Predicted HOI complement ΔpHi gene expression data in identifying highly networked transcription factors

A dataset of human TFs [46] was combined with the set of genes whose transcription was monitored when pHi was raised or lowered [29] (Fig 6A). The overlap of 1,074 TFs was reduced to 689 when including only those TFs expressed at log(CPM) ≥ 2 in cells without pH challenge, a threshold separating the bimodal distribution of expression (Fig 6B) [101]. A further reduction to 224 TFs excluded those with ZF domains. A combined set of TF-target gene relationships was constructed using TFLink, TRRUST, and ARCHS4 data [47–49]. The 224 TFs belonged to 47,178 TF-target pairs, including 6,515 unique targets, with log(CPM) ≥ 2. For each of pHi acidification (with EIPA) or alkalinisation (with NH4Cl) [29], subsets of targets were made for FDR < 0.001 (LOW) and FDR > 0.1 (HIGH), with the region between these thresholds omitted to focus on clearly significant or insignificant fold-change upon ΔpHi. Numbers of (redundant) target genes in these 4 subsets were 19,405 (acid-LOW), 15,968 (acid-HIGH), 4,198 (alk-LOW), and 32,535 (alk-HIGH). Acid pH shift gave a far greater number of targets with significant fold change than alkali pH shift [29]. For non-redundant target genes, the numbers are 2,713 (acid-LOW) and 609 (alk-LOW), with an overlap of 443 target genes.

**Fig 6.**
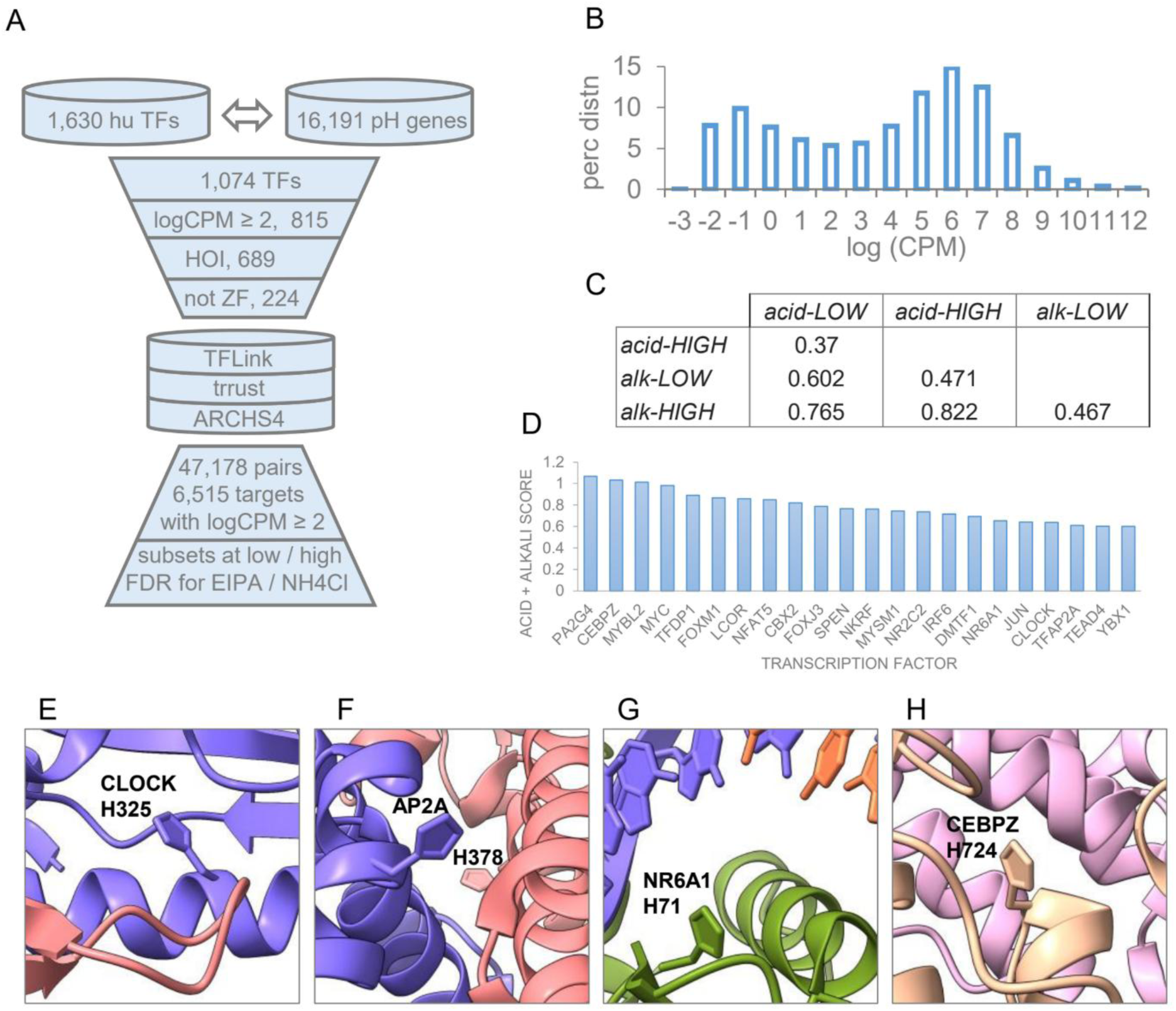
Altered gene expression in pH perturbed cells combined with HOI to identify pH-sensing candidates. (A) Schematic of pipeline to identify TFs common to a dataset of human TFs [46] and a set of genes monitored for altered expression upon pH modulation [29], followed by filtering for significant background expression, inclusion of HOI, and exclusion of ZF domains. TF–target gene pairs were generated from 3 datasets of TF–target relations and filtered for significance of target gene expression. Subsets were formed according to low (0.001) and high (0.1) FDR thresholds for both acidic pH change (EIPA) and alkaline (NH4Cl) [29]. (B) Distribution of log(CPM) values for gene expression in background [29]. (C) Pairwise correlations for 224 (TF) length vectors in acidic/alkaline conditions at low/high FDR, with components equal to the number of targets for each TF in the subset. (D) TFs (22) with the highest scores for contribution to expression changes, combined for acid and alkali changes. Panels (E) to (H) shows 4 examples of HOI amongst the highest scoring TFs. (E) CLOCK-H325 at an interface with BMAL1 (mouse protein, 4f3l [104]). (F) TFAP2A-H378 at a homodimeric interface of the TF, in complex with DNA (8j0k [105]). (G) NR6A1-H71 adjacent to the phosphate backbone of DNA (mouse protein, 5krb [106]). (H) CEBPZ-H724 at a protein-protein interface (from yeast orthologue MAK21-H754 that maps to CEBPZ-H724, 8e5t [107]).

For each of the 4 subsets, 224 length vectors were formed with the number of targets for each TF, and correlations calculated between the vectors (Fig 6C). Correlations are lowest for acid-HIGH with acid-LOW and alk-HIGH with alk-LOW, showing that subsets with significant difference in expression (to unperturbed cells) are distinguishable from subsets without significant differences (as found in the original report [29]). High correlation for acid-HIGH with alk-HIGH and acid-LOW with alk-LOW reflects the overlap of gene expression data for acid and alkali pH changes. Next a score was developed to rank the contribution of each TF to differences between HIGH and LOW FDR subsets. For each of the 224 TFs the number of targets was converted to a percentage of the total number of (redundant) targets in each of the 4 subsets. The difference was calculated between LOW and HIGH subsets, giving values for each of the 224 TFs, with acid or alkali perturbations. The sum of absolute values for acid and alkali gave a single score for each TF. This score, which is agnostic as to whether target gene expression changes are up or down, was ranked. All TFs (22) with a score > 0.6 are shown (Fig 6D).

Applying an additional filter of FDR > 0.05 for TF expression changes focusses on potential pH-dependence of a TF-target pair [29]. For the 22 TFs, this yields CEBPz, LCOR, SPEN, MYSM1, NR6A1, CLOCK, and TFAP2A/AP2A. Searching the RCSB for these 7 human TFs (and orthologues) gives 4 examples, shown with HOI (Fig 6E-H). CLOCK-H325 lies at the interface with TF partner BMAL1 in a complex that is a key element of the mammalian circadian clock (Fig 6E). Circadian rhythms are reported to be altered by an acidic tumour microenvironment [102], which could in principle also change intracellular pH, although the mechanism of coupling acidic pHe and the circadian clock is unclear. TFAP2A-H378 is at a homodimeric interface of the TF, in a complex with DNA (Fig 6F), with roles in development, differentiation and apoptosis. NR6A1 is a nuclear receptor, also involved in development. HOI H71 is adjacent to the phosphate backbone in a complex with DNA (Fig 6G, mouse protein). CEBPZ, a DNA binding factor involved in stress response, is also a component of ribosome assembly complexes. CEBPZ-H724 lies at a protein-protein interface in one such complex for the yeast orthologue MAK21, where H754 maps to H724 (Fig 6H). Again, there is evidence of a link between acidic pHe and the intracellular process of interest (ribosome biogenesis) [103].

### 3.7 Surface HOI associate with flatter regions and, in a subset of complex structures, are mostly predicted to be pH-sensitive at physiological pH

To investigate predicted HOI that are accessible (SASA ≥ 20 Å^2^) in AlphaFold protomers, all proteins with HOI but are not enzymes, transporters, are not annotated to bind ions or contain ZF domains, and which do not have buried HOI (SASA < 20 Å^2^) were selected, giving 1,655 proteins with 2,368 HOI. This subset of proteins has 18,134 histidines that are not predicted HOI, with SASA ≥ 20 Å^2^. Accessible HOI are generally lower in SASA than accessible non-HOI His (Fig 7A), indicative of less protruding, flatter locations that have been associated with interfaces [108].

**Fig 7.**
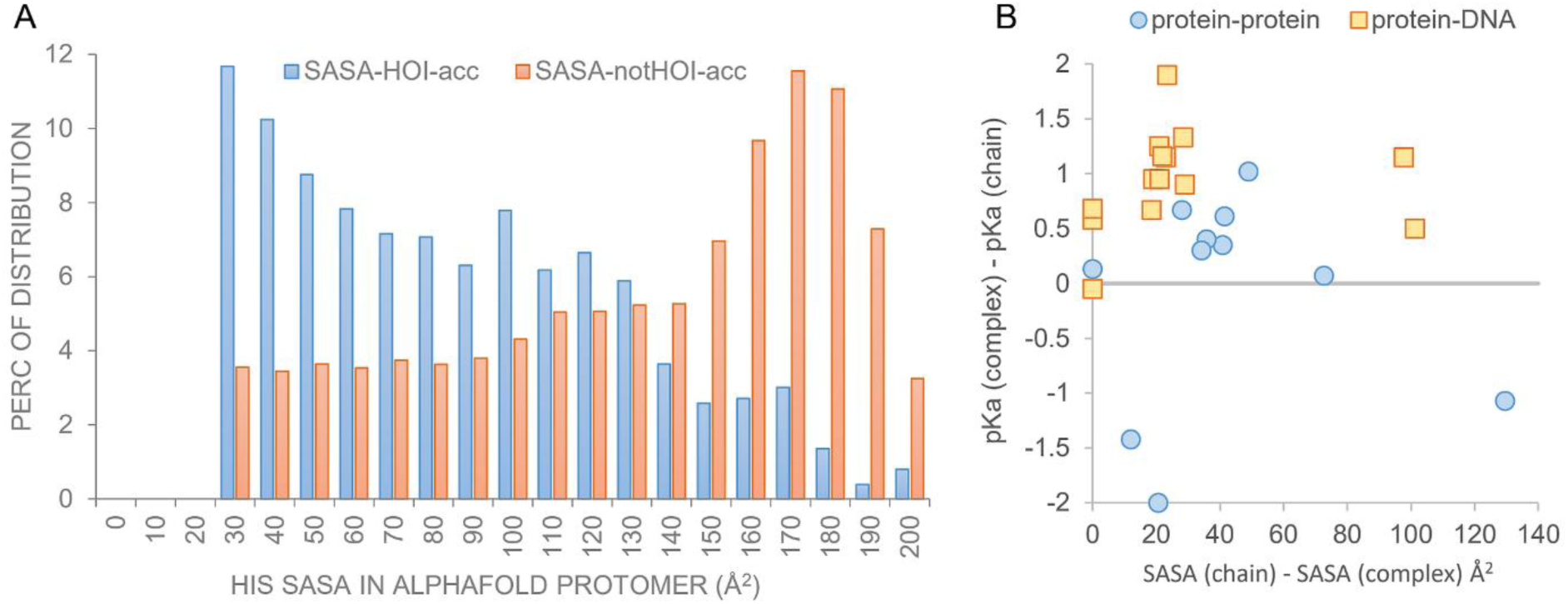
Solvent accessible HOI are generally less exposed than other protomer surface histidines, with significant pKa changes in a subset of complexes. (A) For predicted solvent accessible HOI in 1,655 human proteins that are not enzymes or transporters or ion binding, and do not contain ZF domains or buried HOI, SASA is compared with non-HOI histidines (also with SASA ≥ 20 Å^2^) from the same proteins. (B) Predicted pKa change upon complexation (positive is stabilising for ionised His) are shown for HOI in the 16 systems highlighted with molecular graphics in the current work. Protein-DNA interfaces are differentiated from protein-protein interfaces, and pKa change plotted against SASA change (upon complexation) for each HOI.

Sixteen systems (with PPIs and protein-DNA interfaces) highlight HOI in molecular graphics images in this work. Predicted pKas were computed for those HOI in complex and in free protein extracted from the complex, with the same method used for the FOXC2-DNA complex (Fig 4, relative dielectric of 20 for non-solvent regions). More than 16 HOI are in the calculations due to inclusion of pairs in TF-DNA complexes and independent HOI in the same system. For protein-DNA complexes, one HOI has no SASA or pKa change upon complex formation (Fig 7B). All others have predicted pKa changes that would stabilise ionised HOI and could give pH-dependence around physiological pH. In two protein-DNA interfaces substantial pKa change is predicted without SASA change, showing the role of phosphate backbone in modulating protein pKas (Fig 4C). For HOI at PPIs, the majority have predicted stabilisation of ionised HOI in the complex, although most to a lower degree than for the protein-DNA interactions. Three PPI systems have predicted destabilisation of ionised HOI, a common feature for buried histidines in proteins.

The extent to which interfacial HOI could contribute to functional pH-dependence depends on the overlap of the complexed and uncomplexed pKas with physiological pH. Even with pHi reduction, for example through the cell cycle [88], HOI pKas would need to approach upward from a model compound or intrinsic value (pKa_int) towards 7.0 to be relevant [29]. Histidine pKa_int in the current work is 6.3 [40], compared with 6.5 for two commonly used methods MCCE [109] and PROPKA [110]. Measured pKa for histidine as part of a tripeptide has been reported at 6.8 [111]. The pkAD database curates experimental pKa data across species [112]. The distribution for about 200 histidines is trimodal, with low pKa < 5.5, likely buried His, high pKa > 7.5, likely of functional importance, and a broader mid-range mode that peaks at 6.6 (not shown). Mid-range His pKa (close to 6.6) are unlikely to be charged in normal physiological conditions and could therefore be an estimate of pKa_int. For the calculations of Fig 7B, with pK_int = 6.3, the average of the higher predicted pKa (of free and complexed protein) is 6.9 for protein-DNA interfaces and 6.6 for PPIs. With a model compound pKa of 6.6 these numbers become 7.2 and 6.9, proximal to the physiological pHi range. Functional pH-dependence may be relevant at lower pH values for extracellular proteins, of which there are 114 labelled as secreted in the 1,655 protein set, but none in the 16 structural examples.

Prediction of histidine pKas tend to be more difficult than other pKas [23]. An overall ΔpKa prediction accuracy of around 1 [23] is reducing with ML methods [24], but remains no smaller than the physiological ΔpHi range (about 0.5) of most interest. This is a major incentive to develop alternative protocols for predicting pH-dependent centres, alongside the absence of a complete dictionary of models for PPIs and protein-DNA interactions in human cells, although that area is developing rapidly [21].

## 4. Conclusion

Although much is known about proteins responsible at an overall level for pH-homeostasis within the human cell and its organelles [113], the detail of molecular responses to small ΔpHi and the networks involved, remain largely uncharacterised. Expression changes in response to pHi modulation are extensive [29] and certain families of TFs have conserved histidines, some which are reported to modulate DNA binding at physiological pH [17, 76]. Prediction of His that may be pH-sensing is developed using scores for sequence and structural context [26, 27] rather than pKa calculations, thereby avoiding issues with the accuracy of ΔpKa prediction and with assessment of reliability in the fast-moving field of proteome-wide modelling for protein-protein and protein-DNA complexes. The scores presented here mostly reproduce known pH-sensors, including nuclear receptor and other TF families [17, 76]. Amongst the few test systems missed are histidines in a BRD4 IDR. It is likely that here the evolutionary focus is on function distributed through several pH-sensitive AAs, leading to a requirement for an approach less centred on single sites. Outside of IDRs there are cases, especially in paralogue families, where a histidine just misses the HOI threshold. Histidine can be conserved for functional reasons other than pH-sensing, for example involvement in a specific hydrogen-bond network. Some categories of histidine enrichment were excluded in much of the current analysis, enzyme, transporters, ZF domains, ion-binding. These merit further work using sequence and structural scores. For example, how do His that mediate proton transfer in enzymes compare with other active site His, or can His that are directly involved in proton transport be separated from those that only modulate pH-dependence, in transporters. Separation of HOI inside and outside of ZF domains is also part of future work, with the substantial complication of possible coupled proton and zinc ion binding [114, 115]. Other AAs can be involved in pH-dependence [116]. Inclusion of Asp and Glu in the model requires development of a measure that separates purely salt-bridging AAs from those involved in pH-sensing.

The prominence of DNA-binding proteins in enrichment for HOI illustrated extends to pioneer transcription factors and complexes involved in regulating gene expression and cell division, consistent with the large-scale expression changes reported in response to pHi modulation [28, 29]. Histidines in a subset of protein-DNA complexes are predicted to stabilise complex formation at lower pHi, with ionised His (Fig 7B). This does not necessarily imply a uniform monotonic response to pH, since DNA binding proteins can act to activate or repress transcription [117] with further complexity for factors that tune the balance between activation and repression according to binding affinity [118]. Some CDK-cyclin pairs are prominent in the HOI analysis, which could aid interpretation of data showing that pHi and transit through the cell cycle are coupled [28, 88, 119]. Other areas highlighted in this work include the circadian clock, developmental processes, and ribosome assembly. Further work should include analysis of pathways linked to the TF networks and include extracellular proteins, that experience larger pH changes, for example during inflammation or in the tumour micro-environment [120].

A key question is how to test the predicted scale and detail of pH-dependence that follow from the current analysis. One answer lies in further studies of transcriptome [28, 29] responses to pH change, and in recent parallel analyses of proteome responses. A report of intracellular acidification and regulation of ERK3 degradation includes wider analysis of pHi change on the proteome [121]. Data acquired from modulation of pHe ([122] for example) is relevant for extracellular protein pH-sensing and the downstream intracellular effects. Of particular interest would be large-scale pHi-dependent sets of protein-protein and protein-DNA interactions. For PPIs, a measure of protein stability may be sufficient to highlight pH-sensitive complexes, for example using large-scale solvent-based denaturation of proteins inside cells [123]. Measurement of chromatin opening and TF binding have commonly been made for pertubations such as the addition of inhibitor ([124] for example). Substituting ΔpHi as the perturbation would be a clear strategy.

Experimental investigation of pH-sensing in cells will continue to leverage high throughout methodologies, and predictions are likely to extend the use of machine learning in combining sequence, structural and other data. Although this report does not develop ML protocols, they are indirectly used *via* the key AlphaMissense and AlphaFold components. The most important factor in determining how investigation of pH-sensing in human cells proceeds is to establish whether it is as ubiquitous as predicted in the current work and indicated in earlier reports ([28, 29]).

## Acknowledgements

The authors thank Sifan Zhang for valuable discussions, and staff at the University of Manchester Computational Shared Facility for facilitating storage and processing of data.

## Supporting Information

**S1 Fig.**
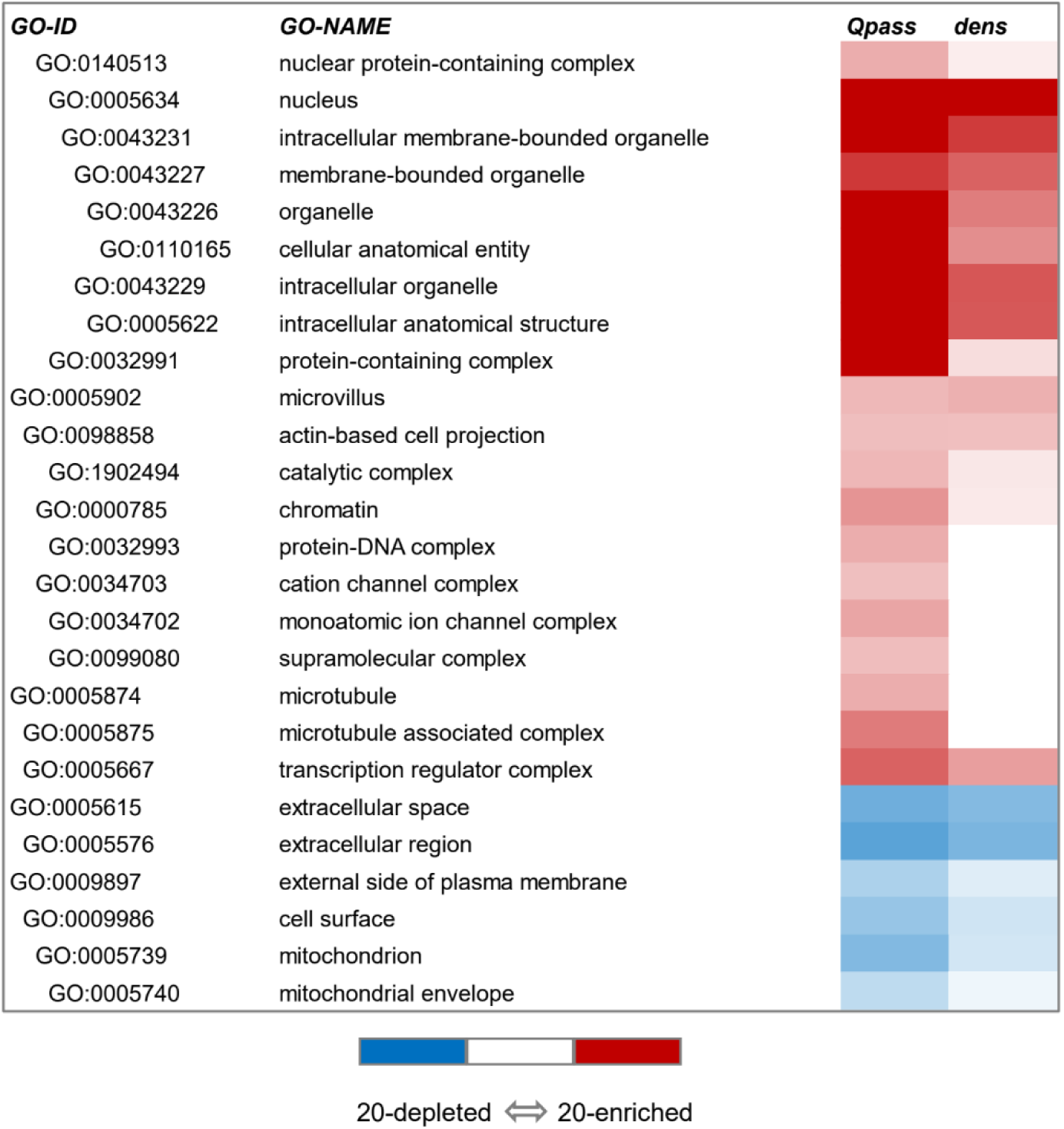
GO Slim component enrichment analysis for glutamine. Heat map of enrichment and depletion for glutamine AAs that pass the same MA and AM thresholds as do predicted HOI (Fig 3).

